# Interkinetic nuclear migration in the zebra1sh retina as a diffusive process

**DOI:** 10.1101/570606

**Authors:** Afnan Azizi, Anne Herrmann, Yinan Wan, Salvador J. R. P. Buse, Philipp J. Keller, Raymond E. Goldstein, William A. Harris

**Author notes:** For correspondence (WAH); (REG). These authors contributed equally to this work (Afnan Azizi, Anne Herrmann).

## Abstract

A major hallmark of neural development is the oscillatory movement of nuclei between the apical and basal surfaces of the neuroepithelium during the process of interkinetic nuclear migration (IKNM). Here, we employ long-term, rapid lightsheet and two-photon imaging of Zebrafish retinas *in vivo* during early development to uncover the physical processes that govern the behavior of nuclei during IKNM. These images allow the capture of reliable tracks of nuclear movements and division during early retinogenesis for many tightly packed nuclei. These tracks are then used to create and test a theory of retinal IKNM as a diffusive process across a nuclear concentration gradient generated by the addition of new nuclei at the apical surface. The analysis reveals the role of nuclear packing at the apical surface on the migration dynamics of nuclei, provides a robust quantitative explanation for the distribution of nuclei across the retina, and may have implications for stochastic fate choice in this system.

## Introduction

The vertebrate nervous system arises from a pseudostratified epithelium within which elongated proliferating cells contact both the apical and basal surfaces. Within these cells, striking nuclear movements take place during the proliferative phase of neural development. More than 80 years ago, these movements, termed interkinetic nuclear migration (IKNM), were shown to occur in synchrony with their cell cycle (***Sauer, 1935***). Under normal conditions, nuclei of proliferating cells undergo mitosis (M) exclusively at the apical surface. During the first gap phase (G1) of the cell cycle, nuclei migrate away from this surface to reach more basal positions by S-phase, when DNA is replicated. In the second gap phase (G2), nuclei migrate rapidly toward the apical surface where they divide again (***Del Bene, 2011; Sauer, 1935; Baye and Link, 2007; Leung et al., 2011; Kosodo et al., 2011; Norden et al., 2009***). The molecular mechanisms that drive the rapid nuclear movement in G2 have been investigated in a number of tissues (***Norden, 2017***). For instance, in the mammalian cortex they are thought to involve microtubules as well as various microtubule motors and actomyosin (***Xie et al., 2007; Tsai et al., 2007***). In the Zebrafish retina, it appears to be the actomyosin complex alone that moves the nuclei to the apical surface during G2 (***Norden et al., 2009; Leung et al., 2011***). In contrast, the nuclear movements during the majority of the cell cycle, in G1 and S phases, have been less thoroughly examined. Although similar molecular motors have been implicated (***Schenk et al., 2009; Tsai et al., 2010***), the underlying processes remain elusive.

Importantly, IKNM is known to affect morphogenesis and cell differentiation in neural tissues (***Spear and Erickson, 2012***), as retinas with perturbed IKNM are known to develop prematurely and to display abnormalities in cell composition (***Del Bene et al., 2008***). Given this regulatory involvement of IKNM in retinal cell differentiation, a deeper understanding of the nuclear movements remains a major prerequisite for insights into the development of neural systems. On a phenomenological level, the movements of nuclei during the G1 and S phases have been shown to resemble a stochastic process in the Zebrafish retina (***Norden et al., 2009; Leung et al., 2011***). During these periods, individual nuclei switch between apical and basal movements at random intervals, leading to considerable variability in the maximum basal position they reach during each cell cycle (***Baye and Link, 2007***). Similarly, in the mammalian cerebral cortex, the considerable internuclear variability in IKNM leads to nuclear positions scattered throughout the entire neuroepithelium in S-phase (***Sidman et al., 1959; Kosodo et al., 2011***). The high variability in the movements of nuclei during G1 and S phases of the cell cycle suggests that passive, rather than active, molecular processes are a main driver of basal migration. This hypothesis was supported by experiments demonstrating similarly variable basalward-biased migration of nuclear-sized microbeads inserted in between cells during IKNM in the mouse cortex (***Kosodo et al., 2011***). Various possible explanations for these passive processes have been put forward. These suggestions include the possibility of direct energy transfer from rapidly moving G2 nuclei (***Norden et al., 2009***), as well as nuclear movements caused by apical crowding (***Kosodo et al., 2011; Okamoto et al., 2013***). Here, we present experiments to test these hypotheses.

Our work relies on the tracks of closely packed nuclei of Zebrafish retinal progenitor cells. The retina of the oviparous Zebrafish is easily accessible to light microscopy throughout embryonic development (***Avanesov and Malicki, 2010***) and has been used for several studies of the movements of nuclei during IKNM (***Baye and Link, 2007; Del Bene et al., 2008; Norden et al., 2009***; ***Sugiyama*** et al., ***2009***; ***Leung et al., 2011***). We find evidence for IKNM being driven by apical crowding and therefore further develop this idea into a mathematical model. Given the seemingly stochastic nature of individual nuclear trajectories, we base the model on a comparison between IKNM and a simple diffusion process. The model reveals the remarkable and largely overlooked importance of simple physical constraints imposed by the overall tissue architecture and allows us to describe accurately the global distribution of nuclei as a function of time within the retinal tissue. In this way, we describe IKNM as a tissue-wide rather than a single-cell phenomenon. In the future, this description might shed light on other aspects of progenitor cell biology, such as cell cycle exit and fate.

## Results

### Generating image sets with high temporal resolution

We imaged fluorescently-labeled nuclei of whole retinas of developing Zebrafish at 2 min intervals, an optimal time period given the difficulty to track nuclei accurately over long times and the increased photobleaching with shorter intervals. We compared movies of retinas imaged at 2 min and at 20 s intervals over a period of 2 hours and found that the improvement in temporal resolution made no difference to our analyses. This suggests that it is unlikely that within each 2 min interval there were important intervening movements that might complicate the analysis.

To follow the nuclei of all cells within a portion of the retina we used H2B-GFP transgenic lines with GFP expression exclusively in the nuclei (Figure 1A). In order to achieve the desired temporal resolution without sacrificing image quality, fluorescence bleaching and sample drift must be minimized as much as possible. The retinas of H2B-GFP embryos were imaged using either a single-angle lightsheet microscope (see Figure 1B for a schematic) or an upright two-photon scanning microscope. Both of these methods yield images with minimal bleaching compared to other microscopic techniques (***Svoboda and Yasuda, 2006; Stelzer, 2015***). However, while the single-angle lightsheet can generate large stacks of images, it is very sensitive to lateral drift due to a small area of high resolution imaging. Therefore, some datasets were produced using two-photon microscopy, which, despite the limitations of scanning time, could produce areas of high resolution images of sufficient size.

**Figure 1.**
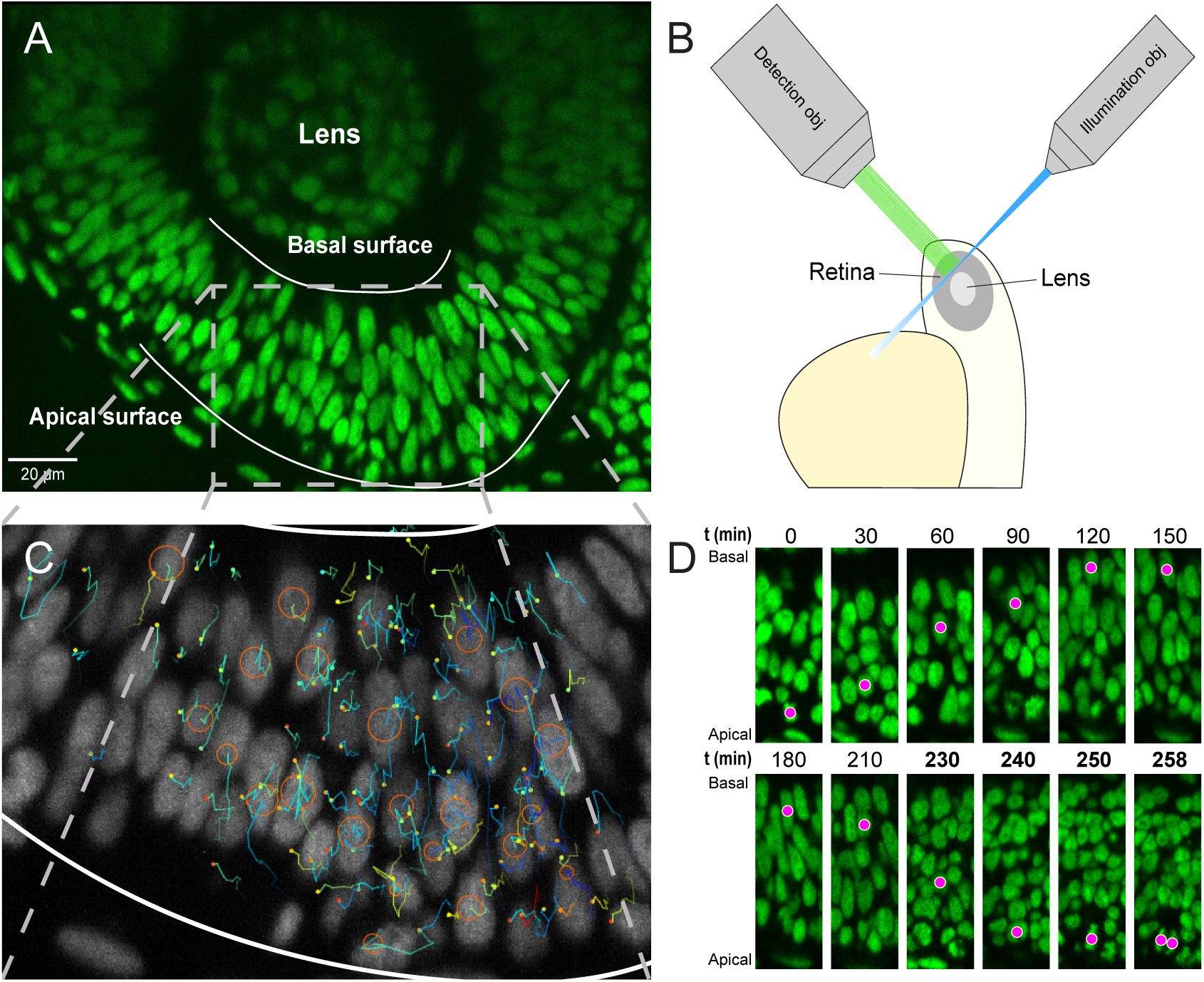
Imaging and tracking fluorescently labeled nuclei. **(A)** A transgenic H2B-GFP embryonic retina imaged using lightsheet microscopy at ∼30 hpf. The lens, as well as apical and basal surfaces are indicated. **(B)** A schematic representation of single-angle lightsheet imaging of the retina. Laser light is focused into a sheet of light by the illumination objective and scans the retina. Fluorescent light is then collected by the perpendicular detection objective. **(C)** Track visualization and curation using the MaMuT plugin of Fiji. All tracks within a region of the retina are curated and visualized. Circles and dots represent centers of nuclei, and lines show their immediate (10 previous steps) track. **(D)** The position of a single nucleus within the retinal tissue from its birth to its eventual division. The magenta dot indicates the nucleus tracked at various time points during its cell cycle. The last 4 panels are at shorter time intervals to highlight the rapid movement of the nucleus prior to mitosis.

Both lightsheet and two-photon microscopes produced images of at least half the retina with a depth of at least 50 µm over several hours in 2 min intervals. The images were processed using a suite of algorithms (***Amat et al., 2015***) to compress them to a lossless format, Keller Lab Block (KLB), correct global and local drift, and normalize signal intensities for further processing. Automated segmentation and tracking of the nuclei were carried out through a previously published computational pipeline that takes advantage of watershed techniques and persistence-based clustering (PBC) agglomeration to create segments and Gaussian mixture models with Bayesian inference to generate tracks of nuclei through time (***Amat et al., 2014, 2015***). Two main parameters greatly affect tracking results, overall background threshold and PBC agglomeration threshold. To obtain best automated tracking results, ground truth tracks were created for a section of the retina over 120 min and were compared to tracks generated over a range of these two parameters. The best combination of the two parameters was chosen as the one with highest tracking fidelity and lowest amount of oversegmentation over that interval.

The most optimal combination of parameters yielded an average linkage accuracy, from each time point to the next, of approximately 65%. Hence, extensive manual curation and correction of tracks were required. Tracking by Gaussian mixture models (TGMM) software generates tracks that can be viewed and modified using the Massive Multi-view Tracker (MaMuT) plugin of the Fiji software (***Wolff et al., 2018; Schindelin et al., 2012***). A region of the retina with the best fluorescence signal was chosen and all tracks within that region were examined and any errors were corrected. The tracks consist of sequentially connected sets of 3D coordinates representing the centers of each nucleus (Figure 1C), with which their movement across the tissue can be mapped over time. For example, Figure 1D shows IKNM of a single nucleus tracked from its birth, at the apical surface of the retina, to its eventual division into two daughter cells.

### Analysis of nuclear tracks

This process yielded tracks for hundreds of nuclei, across various samples, over time intervals of at least 200 min. We used custom-written MATLAB scripts to analyze these tracks. The aggregated tracks of the main dataset, in Cartesian coordinates, for all tracked lineages is shown in Figure 2A. Single tracks for any given time interval can be extracted and analyzed from this collection. In order to transform the Cartesian coordinates of the tracks into an apicobasal coordinate system, we drew contour curves at the apical surface of the retina (e.g. see Figure 1A) separating RPC nuclei from the elongated nuclei of the pigmented epithelium. We then calculated curves of best fit (second degree polynomials) in both the XY and YZ planes. Assuming that the apical cortex is perpendicular to the apicobasal axis of each cell, displacement vectors of the nuclei at each time point can be separated into apicobasal and lateral components. Since, in IKNM, the apicobasal motion is that of interest, we used this component for our remaining analyses.

**Figure 2.**
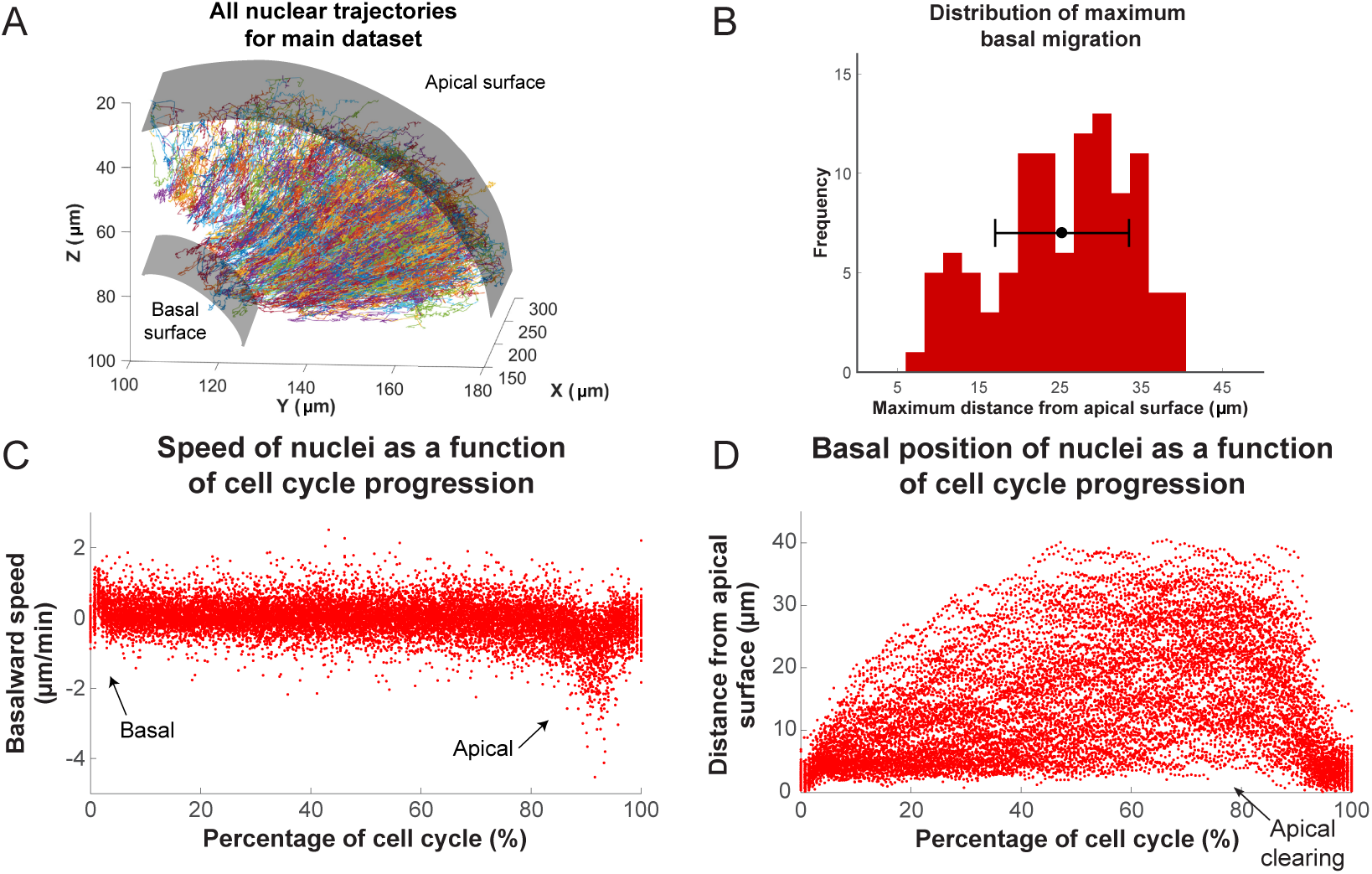
Analysis of nuclear tracks during IKNM. **(A)** Extracted trajectories of nuclei in 3 dimensions. All curated tracks of the main dataset over 400 minutes in the region shown in Figure 1C are presented. **(B)** The distribution of maximum distances reached away from the apical surface by nuclei during their completed cell cycles. The mean and one standard deviation are shown. **(C)** The speed distribution of nuclei over complete cell cycles. The cell cycle lengths of all nuclei were normalized and superimposed to highlight the early basal burst of speed, as well as pre-division apical rapid migration. The speeds between these two periods are normally distributed. **(D)** Position of nuclei as measured by their distance from the apical surface over normalized cell cycle time. Even though all nuclei start and end their cell cycle near the apical surface, they move out across the retina to take positions in all available spaces, creating an apical clearing as indicated.

Figure 2C,D shows the speed and position of tracked nuclei of the same dataset, over the duration of their cell cycle, for all cells that went through a full cell cycle. While all nuclei behave similarly minutes after their birth (early G1) and before their division (G2), their speed of movement and displacement is highly variable for the majority of the time that they spend in the cell cycle (Figure 2C,D). Most daughter nuclei move away from the apical surface, within minutes from being born, with a clear basalward bias in their speed distribution (Figure 2C). This abrupt basal motion of newly divided nuclei has also been recently observed by others (***Shinoda et al., 2018; Barrasso et al., 2018***). However, immediately after this brief period, nuclear speeds become much more equally distributed between basalward and apicalward, with a mean value near 0. Such a distribution is indicative of random, stochastic motion, which in turn leads to a large variability in the position of nuclei within the tissue (away from the apical surface) during the cell cycle (Figure 2B).

Interestingly, except during mitosis, we find an apical clearing of a few microns for dividing cells (Figure 2D). We checked to see if this was an artifact of measuring the distance to nuclear centers due to nuclear shape, as nuclei are rounded during M phase but are more elongated along the apicobasal axis at other times. We found no significant difference between average length of nuclear long axis when measured for nuclei right before their division compared to nuclei chosen randomly from any other time point within the cell cycle, indicating that this clearing is likely to have a biological explanation, such as the preferential occupancy of M phase nuclei to the apical surface during IKNM.

### Basal movement of nuclei is driven like a diffusive process

Previous work has shown that when RPCs are pharmacologically inhibited from replicating their DNA, their nuclei neither enter G2 nor exhibit rapid persistent apical migration that normally occurs during the G2 phase of the cell cycle (***Leung et al., 2011; Kosodo et al., 2011***). A more surprising result of these experiments is that the stochastic movements of nuclei in G1 and S phases also slow down considerably during such treatment (***Leung et al., 2011***). It was, therefore, suspected that the migration of nuclei of cells in G2 toward the apical surface jostles those in other phases (***Norden*** et al., ***2009***). We therefore searched our tracks for evidence of such direct kinetic interactions among nuclei by correlating the speed and direction of movement of single nuclei with their nearest neighbors. These neighbors were chosen such that their centers fell within a cylindrical volume of a height and base diameter twice the length of long and short axes, respectively, of an average nucleus. Figure 3A shows the lack of correlation between the speed of movement of nuclei and the average speed of their neighbors. We further categorized the neighboring nuclei by their position in relation to the nucleus of interest (along the apicobasal axis), their direction of movement, and whether they were moving in the same direction of the nucleus of interest or not. None of the resulting eight categories of neighboring nuclei showed a correlation in their average speed with the speed of the nucleus of interest. Furthermore, we considered the movement of neighboring nuclei one time point (2 min) before or one time point after the movement of the nucleus of interest. Yet, we still found no correlation between these time-delayed and original speeds. These results suggest that there does not appear to be much transfer of kinetic energy between neighboring nuclei.

**Figure 3.**
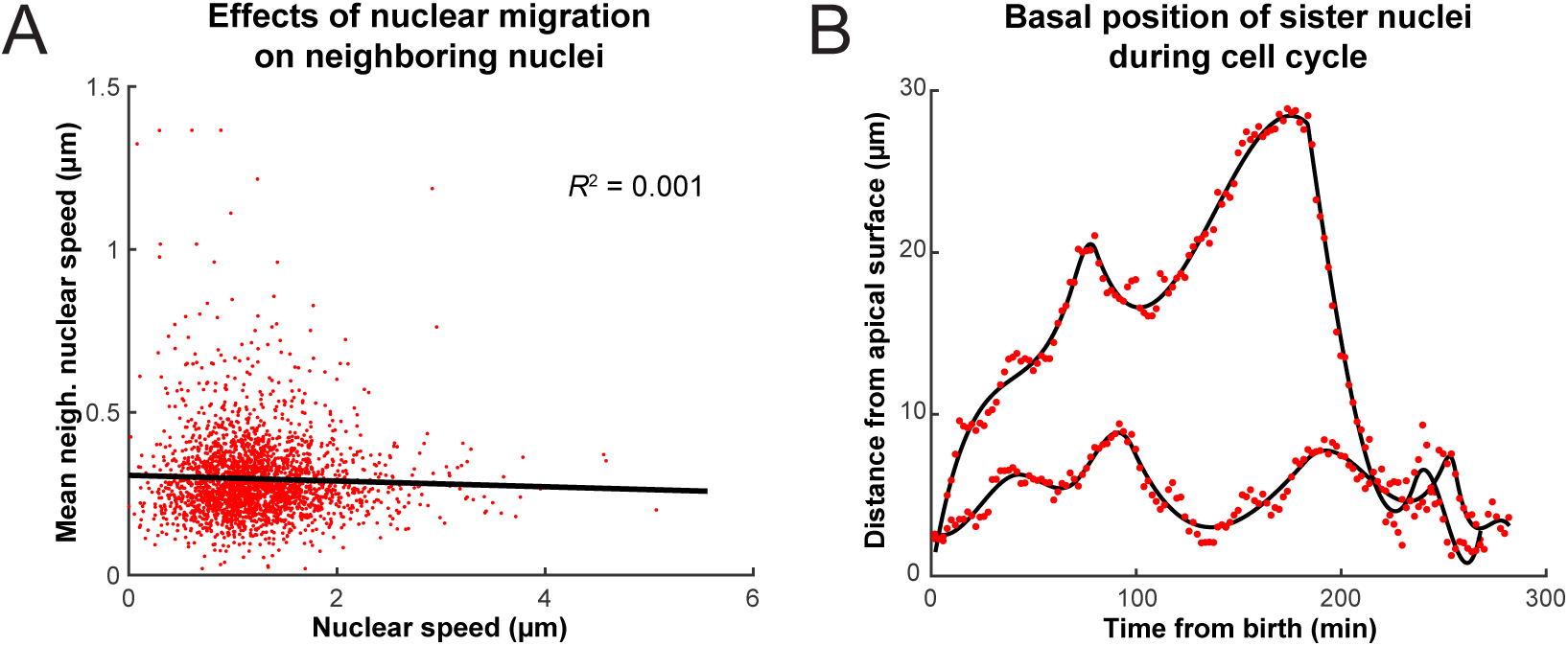
**(A)** Average speed of nuclei neighboring a nucleus of interest as a function of the speed of that nucleus. **(B)** The positions of two sister nuclei at each time point imaged (red circles) over their complete cell cycle. The black lines are spline curves indicating the general trend of their movements.

Another hypothesis advanced for variability in basal IKNM is that the nuclear movements are driven by apical crowding (***Kosodo et al., 2011; Okamoto et al., 2013***). How apical crowding might result in basal IKNM can be understood by comparing IKNM to a diffusive process. In diffusion, a concentration gradient drives the average movement of particles from areas of high to areas of low concentration. However, despite the average movement being directed, each individual particle’s trajectory is a random walk (***Reif, 1965***). Similarly, during IKNM a gradient in nuclear concentration is generated because nuclei divide exclusively at the apical surface. If basal IKNM were comparable to diffusion, this nuclear concentration gradient would be expected to result in a net movement of nuclei away from the area of high nuclear crowding at the apical side of the neuroepithelium (***Miyata et al., 2015; Okamoto et al., 2013***). Indeed, in IKNM we find that each individual nucleus’ trajectory resembles a random walk (***Norden et al., 2009***). Therefore, for the cells in the G1 and S phases (which account for more than 90% of the cell cycle time in our system), IKNM has, at least on a phenomenological level, the main features of a diffusive process.

To test further whether we can indeed describe IKNM using a model of diffusion, we first asked what would happen to the concentration gradient if we blocked the cell cycle in S phase, which inhibits both the apical movement of the nuclei in G2 and mitosis at the apical surface. If the comparison to diffusion were valid, we expect the blockage to abolish the build-up and maintenance of the concentration gradient. We, therefore, compared the normally evolving distribution of nuclei in control retinas with those measured from retinas where the cell cycle was arrested at S-phase using a combination of hydroxyurea (HU) and aphidicolin (AC) (***Norden et al., 2009; Icha et al., 2016***). We counted the number of nuclei in a three dimensional section of the retina containing approximately 100 nuclei, at equal time intervals, starting with 120 min after drug treatment. The delay ensured that almost all cell divisions, from nuclei that had already completed the S phase at the time of treatment, had taken place. As expected from the diffusion model (Figure 4D), over the course of 160 min, the mean of the nuclear distribution moved further towards the basal surface in treated retinas, and the concentration difference between the apical and basal surfaces diminished (Figure 4A,C). In contrast, in control retinas the mean of the nuclear distribution moved towards the apical surface (Figure 4B,C) as the gradient continued to build up. Hence, these results support the suitability of a diffusive model to describe the basal nuclear migration during IKNM.

**Figure 4.**
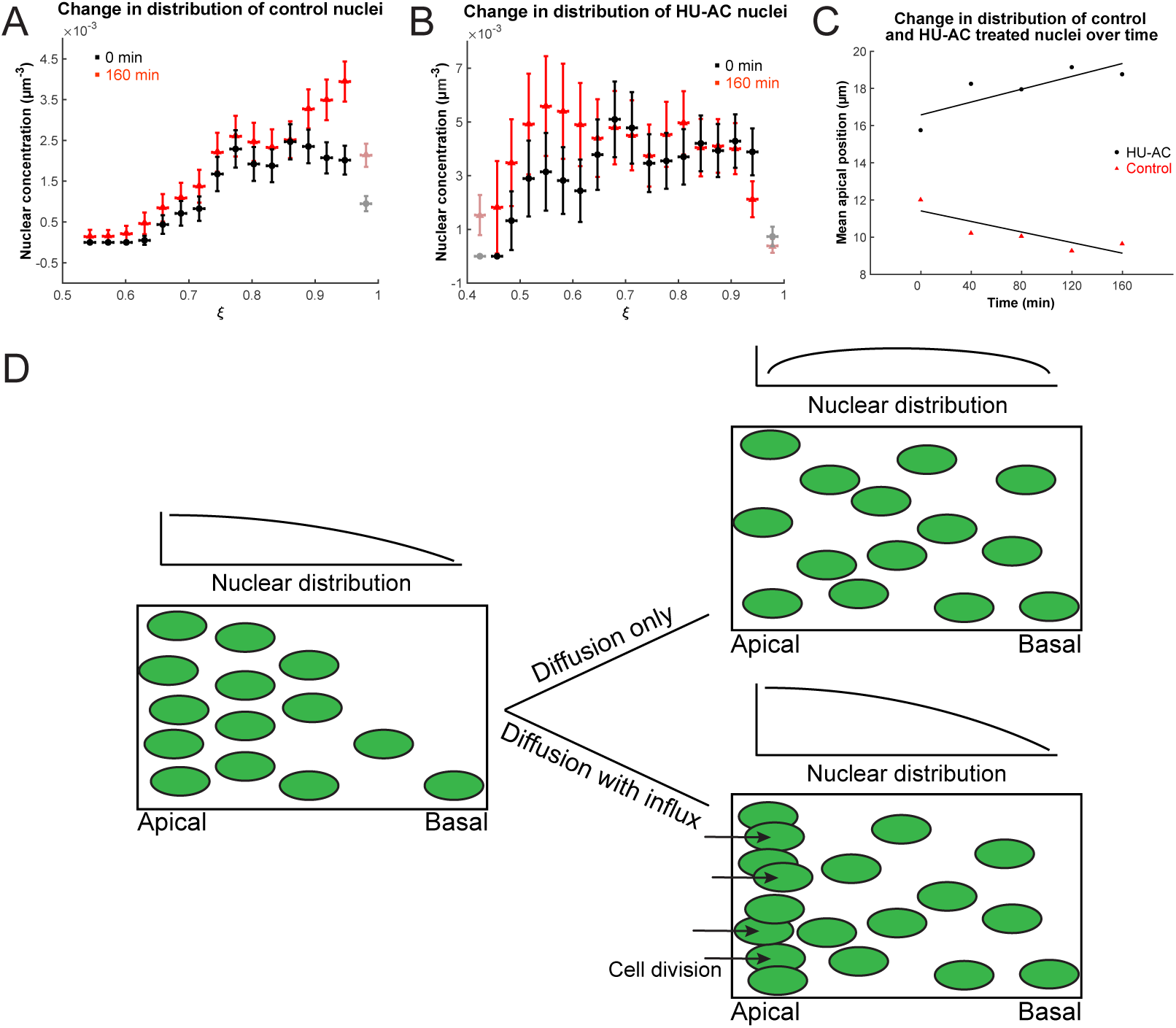
Nuclear concentration gradient across the apicobasal axis of the retina. The concentration of nuclei is higher near the apical surface compared to the basal surface. **(A)** In the control retina the nuclear concentration gradient builds up over time. **(B)** Blocking apical migration and division of nuclei, by inhibiting S phase progression, leads to a shift in the distribution of nuclei towards the basal surface in the HU-AC treated retina. **(C)** The shift in the distribution of nuclei under HU-AC treatment when compared to the untreated retina. The number of nuclei away from the apical surface increases consistently over time in the absence of cell division, but remains the same when new nuclei are constantly added at the apical surface. **(D)** A schematic of how a diffusion model would work in the context of IKNM in the retina. A concentration gradient of nuclei (left) would drive the net movement of nuclei from the apical surface to the basal surface. However, without maintenance of the gradient, the drive for this net migration is lost (top right). In the retina, the gradient is maintained through cell divisions at the apical surface, modeled as a one way influx across the apical surface (bottom right), continuously driving the net movement basally.

### An analytical diffusion model of IKNM

To investigate whether a diffusion model would also provide a useful quantitative description for IKNM, we formalized the process of IKNM in mathematical terms. This formalization again focuses on the crowding of nuclei at the apical side of the tissue. Crowding can be thought of, in mathematical terms, as creating a gradient in nuclear concentration *c* along the apicobasal direction of the retina. In contrast, we assumed no dependence of the nuclear concentration on the lateral position within the tissue. Thus we employed the diffusion equation for the nuclear concentration *c*(*r, t*) as a function only of the apicobasal distance *r* and time *t*. The retina can be approximated as one half of a spherical shell around the lens, and thus we use spherical polar coordinates with the origin of the coordinate system at the center of the lens, the basal surface at *r* = *b* and the apical surface at *r* = *a* (Figure 5B). We first study the simplest diffusion equation for this system, in which there is a diffusion constant *D* independent of position, time, and *c* itself, namely

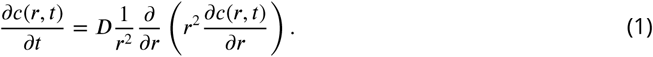

By analyzing the experimental data we seek to determine *D*. This equation provided the basis for our mathematical description of IKNM in terms of a diffusion process.

**Figure 5.**
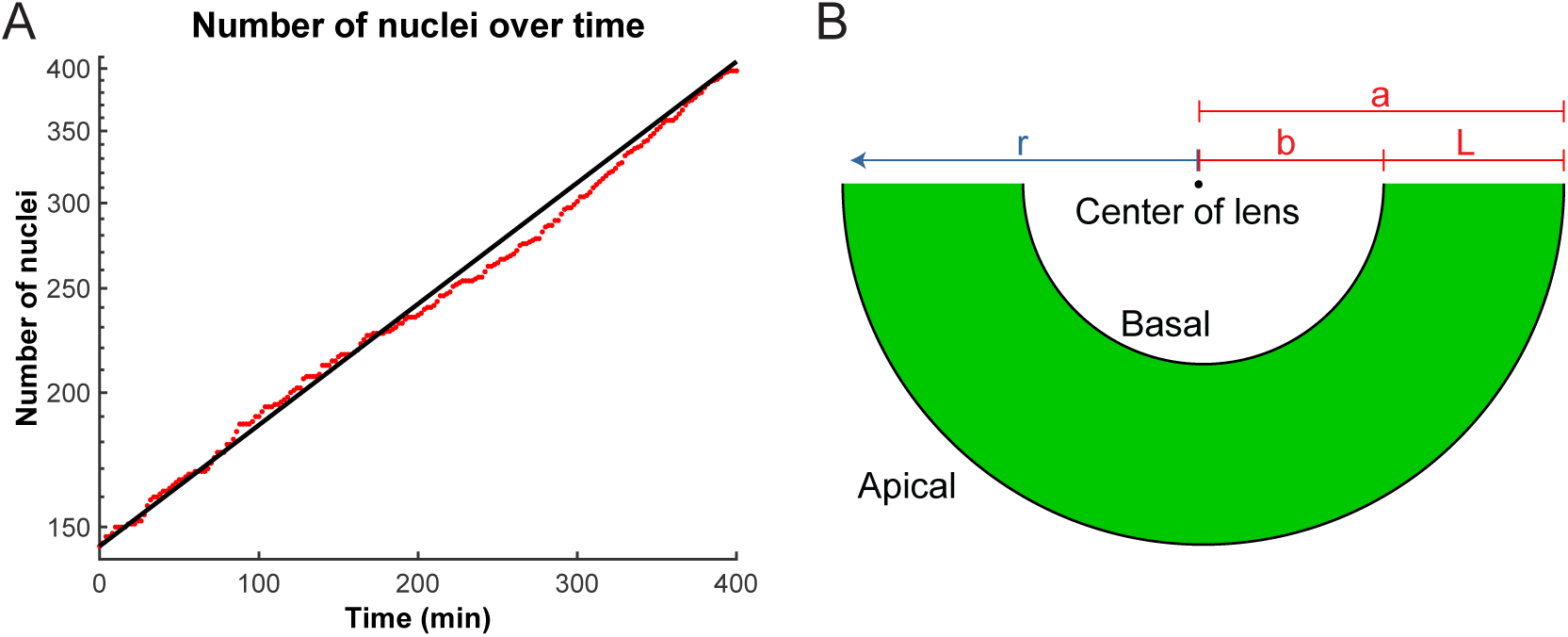
**(A)** Number of nuclei grows exponentially during the proliferative stage of the retinal development. A line can be 1t to the log-lin graph of nuclear numbers as a function of time to extract the doubling time (cell cycle length) in this period. **(B)** A schematic of the retina indicating the variables used in the diffusion model of IKNM. a: distance from center of lens to apical surface; b: distance from center of lens to basal surface; L: thickness of the retina; r: distance from center of lens for each particle.

In addition to Equation 1, we also needed to specify the boundary conditions adequate to describing IKNM. As mentioned above, we focused our description of IKNM on the apical crowding of nuclei. Since nuclei only divide close to the apical surface of the tissue, we treat mitosis as creating an effective influx of nuclei through the apical boundary. To quantify this influx, we extracted the number of cells *N*(*t*) as a function of time. As during the stages of development examined here cells are neither dying nor exiting the cell cycle (***Biehlmaier et al., 2001***), we assumed that the number of cell divisions is always proportional to the number of currently existing cells. This assumption predicts an exponential increase in the number of cells or nuclei, over time, also recently found by ***Matejčić et al.*** (***2018***):

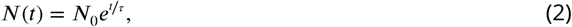

where *N*_0_ is the initial number of nuclei and *r* = *T*_*P*_ / ln 2, with *T*_*P*_ the average cell cycle length. Figure 5A shows the agreement between the theoretically predicted curve *N*(*t*) with the experimentally obtained numbers of nuclei over time. Having obtained *N*_0_ and *T*_*P*_ from our experimental data, the predicted curve does not have any remaining free parameters and thus no fitting is necessary. Thus, the obtained description for the number of nuclei over time, Equation 2, was used to formulate the influx boundary condition for our mathematical model

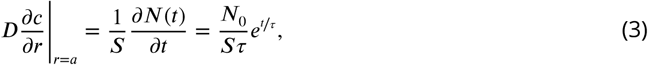

with *S* the apical surface area of our domain of interest. In contrast to the apical side of the tissue, there is no creation (or depletion) of nuclei at the basal side (***Matejčić et al., 2018***), and hence a no-2ux boundary condition,

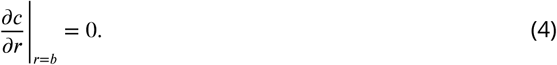

Equations 1, 3 and 4 fully specify this simplest mathematical model of IKNM.

From these equations we can derive an expression for the concentration of nuclei *c*(*r, t*) in the retinal tissue. To this end, we introduced dimensionless variables for space and time,

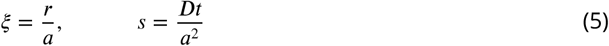

and further define *p* = *b*/*a <* 1. The exact solution for the nuclear concentration, whose detailed derivation is given in the Appendix, is

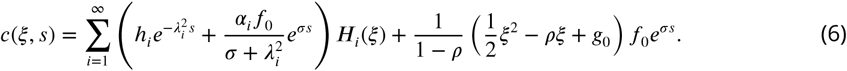

The first terms within parentheses describes the decay over time of the initial condition *c*_exp_(*ξ, s* = 0). Here *λ*_*i*_ are the eigenvalues and *H*_*i*_(*ξ*) the eigenfunctions of the radial diffusion problem, and the coefficients *h*_*i*_ are determined from the experimental initial conditions (see Methods). The second terms within the sum and the final term on the right hand side of Equation 6 are constructed such that the solution fulfills the boundary conditions 3 and 4. In the last term, the constant *g*_0_ was obtained using the constraint that the volume integral of the initial concentration yields the initial number of nuclei *N*_0_. *f*_0_, *σ* and *α*_*i*_ emerge within the calculation of the solution and are specified in the Appendix. Thus, the effective diffusion constant *D* in Equations 1 and 6 is the only unknown in the model.

### The linear model is accurate at early times

As mentioned before, the only parameter in the solution 6 is the effective diffusion constant *D*. To determine this from the data, the experimentally obtained distribution of nuclei in the retinal tissue was first converted into a concentration profile. Then, the optimal *D*-value, henceforth termed *D*\*, was obtained using a minimal-*χ*^2^ approach. The value obtained within the linear model for a binning width of 3 µm and an apical exclusion width of 4 µm is 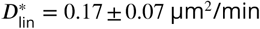. Using this, we can examine the decay times of the different modes in the first term of Equation 6. The slowest decaying modes are the ones with the smallest eigenvalues *λ*_*i*_ and we find that the longest three decay times are *𝒥*_1_ ≈ 1325 min, *𝒥*_2_ ≈ 350 min and *𝒥*_3_ ≈ 158 min. This shows that indeed all three terms of Equation 6 are relevant on the timescale of our experiment and need to be taken into account when calculating the concentration profile. The corresponding plots of *c*(*ξ, s*) are shown in Figure 6A-C. As can be seen from this figure, the diffusion model fits the data very well at early times, *t ≤* 200 min. However, for *t ≥* 200 min the model does not fit the data as well; the experimentally observed nuclear concentration levels off at a value between 4.00 and 4.50 × 10^−3^ µm^−3^ (Figure 6D), an aspect that is not captured by the model of linear diffusion.

**Figure 6.**
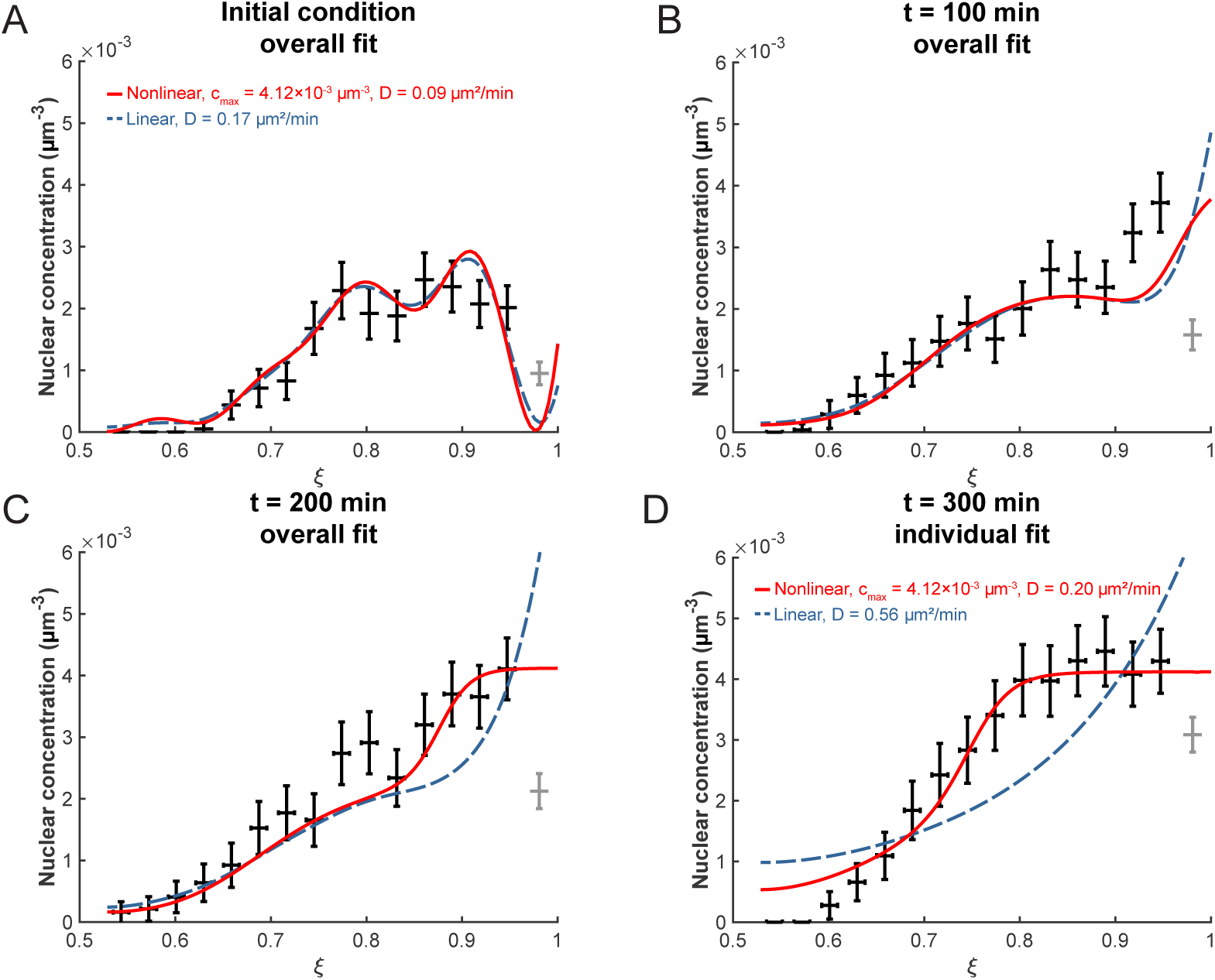
**(A)** The initial experimental concentration profile of nuclei at *t* = 0 min as well as the calculated initial condition curves (see Methods Equation 11) for the linear (red solid line) and nonlinear (blue dashed line) models. B,C,D) The fit of the models to experimental distribution of nuclei after 100 min **(B)**, 200 min **(C)**, and 300 min **(B)** are shown. For the first three graphs, the best fit over all 100 intervening time points were used with the corresponding diffusion constants shown in **(A)**. For t = 300 min, the best fit at that time point only was used with the corresponding diffusion constants indicated.

One particular aspect of the biology that the linear model neglects is the spatial extent of the nuclei. In a linear diffusion model, particles are treated as point-like and non-interacting. However, our microscopy images (see Figure 1A) clearly indicate that the nuclei have finite incompressible volumes, so that their dense arrangement within the retinal tissue would lead to steric interactions once the nuclear concentration is sufficiently high, and moreover that the packing density of nuclei can not exceed a maximum value dictated by their geometry. Next, we examine whether accounting for these effects leads to a more accurate theory.

### Nonlinear extension to the model

If we write the diffusion equation 1 in the form

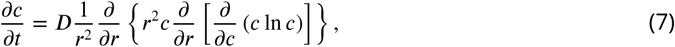

we can identify the term *c* ln *c* as proportional to the entropy *𝒥* of an ideal gas, and its derivative with respect to *c* as a chemical potential. In an ideal gas, all particles are treated as point-like and without mutual interactions. In order to include the spatial extent of particles, we estimate the entropy using the model of a lattice gas, a system in which space is divided into discrete sites which can either by empty or occupied by a single gas particle. Due to the discrete lattice, particles cannot get closer than the lattice spacing from each other, and there is a maximum possible concentration *c*_max_ (***Huang, 1987***). In this system the entropy takes the form

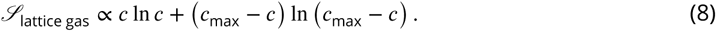

Substituting this expression for the term *c* ln *c* in 7, we obtain the nonlinear diffusion equation

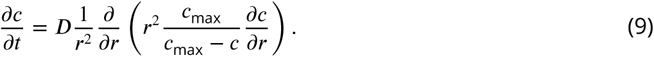

Adjusting the boundary conditions at the apical side accordingly leads to

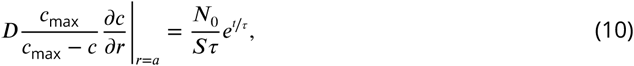

while the basal boundary condition remains the same as Equation 4. Together, Equation 9 and the boundary conditions in Equations 10 and 4 represent an extension to the diffusion model for IKNM, which now accounts for steric interactions between the nuclei. The maximum concentration *c*_max_ incorporated in this model was obtained, as described in the Methods, by considering a range of nuclear radii and the maximum possible packing density for aligned ellipsoids (***Donev et al., 2004***).

Similar to fitting the linear model, we also need to establish a description of the initial condition. To make both models consistent with each other, we employ the linear model’s initial condition, Equation 6 at *s* = 0 with *h*_*i*_ as obtained from Equation 11, as an initial condition for this nonlinear model as well (Figure 6A). The concentration profile in the nonlinear model and its derivative were obtained numerically using the MATLAB pdepe solver. Fitting this concentration profile to the data was again by means of a minimal-*χ*^2^ approach. When the optimization took data points up to *t* = 200 min into account, we find 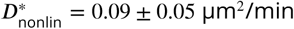 (Figure 6, Table 1). As can be seen, by choosing *c*_max_ correctly, an excellent fit to the data can be obtained. These results show that a lattice-gas based diffusion model is indeed suitable to describe time evolution of the nuclear concentration profile in zebrafish retina tissue during IKNM over several hours of development.

**Table 1.** List of best-fit diffusion constants *D*\*, their standard deviations and probabilities for the studied conditions.

| | $D_{\text{nonlin}}^*$ ($\mu\text{m}^2/\text{min}$) | $\sigma_D$ ($\mu\text{m}^2/\text{min}$) | $P_\chi(\chi^2; \nu)$ |
| --- | --- | --- | --- |
| Normal | 0.09 | 0.05 | 0.49 - 0.51 |
| Normal (repeat sample) | 0.10 | 0.06 | 0.47 - 0.48 |
| High T | 0.13 | 0.08 | 0.42 |
| Low T | 0.06 | 0.05 | 0.69 - 0.7 |

### Incubation temperature has direct effects on IKNM

The diffusion model may also address mechanistic questions about IKNM in retinas growing under varying experimental conditions. Zebrafish embryos are often grown at different temperatures to manipulate their growth rate (***Kimmel et al., 1995; Reider and Connaughton, 2014***), but it has been unclear how the nuclei in the retina behave at these different temperatures. To examine this issue, we grew the embryos at the normal temperature of 28.5 °C overnight and then incubated them at lower temperature (LT) of 25 °C or higher temperature (HT) of 32 °C during imaging. We could directly measure the change in average cell cycle length from experimental data and found that in HT, it is 205.5 min, while in LT, it is a much longer 532.78 min. We were then able to use these values in the model to investigate whether the change in temperature influences the processes that determine the effective diffusion constant of the nuclei. The resulting values for 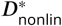 are summarised in Table 1. Based on these values, two-sided *t*-tests (see Methods) confirmed that there is no significant difference between the *D*-values obtained from the two normal condition data sets. In contrast, *D*-values for the LT and HT data sets were significantly different from the normal ones, with *p* ≤ 0.01. These results indicate, that aside from its effect on cell cycle length, incubation temperature is likely to influence IKNM directly by altering the mobility of nuclei, here represented by the effective diffusion constant *D*.

## Discussion

In this work, we have shown that high density nuclear trajectories can be used to tease apart the possible physical processes behind the apparently stochastic movement of nuclei during interkinetic nuclear migration. Firstly, we generated these trajectories using long-term imaging and tracking of nuclei with high spatial and temporal resolution within a 3-dimensional segment of the Zebrafish retina. Analysis of speed and positional distributions of more than a hundred nuclei revealed a large degree of variability in their movements during G1 and S phases. Although this variability had been observed before, previous experiments had only considered sparsely labeled nuclei within an otherwise unlabeled environment (***Baye and Link, 2007; Norden et al., 2009; Leung et al., 2011***). Thus, our results provide an important account of the variability of IKNM on a whole tissue level. In effect, the variability of IKNM means that nuclear trajectories appear stochastic during the majority of the cell cycle. Previously, it had been suggested that the origins of this apparent stochasticity lay in the transfer of kinetic energy between nuclei in G2 exhibiting rapid apical migration to nuclei in G1 and S phases of the cell cycle, much as a person with an empty beer glass may nudge away other customers to get to the bar (***Norden et al., 2009***). However, we found no evidence for direct transfer of kinetic energy between nuclei and their immediate neighbors. Recently ***Shinoda et al.*** (***2018***) have also provided evidence that suggests direct collisions do not contribute to basal IKNM.

Another possibility is that the stochastic trajectories of G1 and S nuclei could be a result of passive displacements, arising from a diffusive process depending on a nuclear concentration gradient between the apical and basal sides of the tissue (***Miyata et al., 2015***). This gradient could be formed by nuclear divisions taking place exclusively at the apical surface. We confirmed the presence of such a gradient by calculating the nuclear concentration along the apicobasal dimension within the retinal tissue at various time points. Further, to probe the source of the gradient, we treated the Zebrafish retina with HU-AC to stop the cell cycle in S phase. While we observed the build-up of the nuclear concentration gradient over time in the control retina, the nuclear distribution flattened when cell division was inhibited with HU-AC treatment. These phenomenological similarities between IKNM and diffusion suggested the diffusive model. This model includes two key features: most importantly, it focuses on the crowding of nuclei at the apical surface of the tissue, here included as the apical boundary condition. Additionally, in the nonlinear extension of the model, it incorporates a maximum possible nuclear concentration. This addition provided a striking overall improvement to the fits to experimental data over periods of many hours. The resulting difference in the obtained *D*-values between the linear and nonlinear versions of our model can be understood heuristically when closely examining the difference between Eqs. 1 and 9. The latter introduces the new term *c*_max_/(*c*_max_ - *c*) which one could think of loosely as corresponding to an effective, concentration dependent diffusion constant 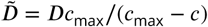. In general 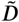 will vary across the tissue thickness and, since *c >* 0 for most of the retinal tissue, 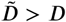. Therefore, averaging across the retina tissue, 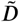 might actually be in very good agreement with the *D*-value found in the linear model. However, the fact the linear model fails to describe, and which leads to a better representation of the data using the nonlinear model, is that the mobility of the nuclei is likely to be concentration dependent.

The underlying processes causing IKNM during the G1 and S phases of the cell cycle in pseudostratified epithelia have been largely elusive. Several partially competing ideas have been put forward, ranging from the active involvement of cytoskeletal transport processes to passive mechanisms of direct energy transfer or movements driven by apical nuclear crowding (***Schenk et al., 2009***; ***Tsai*** et al., 2010; ***Norden et al., 2009; Kosodo et al., 2011***). The fact that inanimate microbeads migrate much like nuclei during IKNM in the mouse cerebral cortex (***Kosodo et al., 2011***) suggests that active, unidirectional intracellular transport mechanisms are not directly responsible for these stochastic movements. Instead, we showed that a passive diffusive process which takes steric interactions between nuclei into account produces an excellent representation of the time evolution of the actual nuclear distribution within the retinal tissue during early development. Consequently, our work builds on earlier models of apical crowding based on *in silico* simulations of IKNM (***Kosodo et al., 2011***). Having said this, it remains to understand the general scale of the diffusion constant (*D* ∼0.1 *µ*^2^/min) from microscopic considerations, perhaps analogous to those used to relate random walks to diffusion (***Goldstein, 2018***). In addition, our work revealed the remarkable importance of simple physical constraints imposed by the overall tissue architecture, which could not be explored in previous studies which tracked sparse nuclei, and thus lacked the means to explore the effect of such 3-dimensional arrangements. Hence, we paid special attention to the spherical shape of the retina and the concentration of nuclei in that space. Examining the evolution in distribution of nuclei over time unveils the importance of spatial restriction due to the curvature of the tissue. Additionally, the size of the nuclei in comparison to the neuroprogenitor cells leads to the emergence of a maximum nuclear concentration which must be taken into account to accurately model IKNM.

By inhibiting cell cycle progression or changing temperature, we used our model to shed some light on some of the properties of and mechanisms of the stochastic movements of nuclei during IKNM. From our results and previous studies, we knew that cell cycle length is affected by change in incubation temperature (***Kimmel et al., 1995; Reider and Connaughton, 2014***). However, our results also indicate a significant influence of temperature on the mobility of nuclei and thus the underlying processes controlling their movement. For example, the speed and dynamic properties of both the microtubule and actomyosin systems are dependent on temperature and could in part explain the changes in the diffusion constant that we see as a function of temperature (***Hartshorne et al., 1972; Hong et al., 2016***) as the diffusion constant may be influenced by stochastic associations with motor proteins or the physical properties of the epithelium. However, a much closer examination of molecular mechanisms driving stochastic nuclear movements is required to better understand the connections between these phenomena, as we are far from understanding the nature of forces involved in this process. Furthermore, the diffusion constant reported here contains all types of nuclear movement during IKNM as it is derived from the changing nuclear concentration profile over time. However, it is not immediately clear what the contribution the rapid apical migration to this overall diffusion constant may be. Nonetheless, despite the large displacement during rapid apical migration at G2, this phase only accounts for about 8% of the cell cycle (***Leung et al., 2011***). Therefore, given this small portion of the cell cycle when rapid migration can happen and the good agreement of our calculated diffusion constant with those previously reported in the literature for individual nuclei (***Leung et al., 2011***), the proposed model appears to describe tissue-wide IKNM quite well.

The physiological consequences of nuclear arrangements and the IKNM movements associated with all pseudostratified epithelia are not well understood. Our results provide a quantitative description of the stochastic distribution of the nuclei across the retina. This distribution has been implicated in stochastic cell fate decision making of progenitor cells during differentiation (***Clark et al., 2012; Baye and Link, 2007; Hiscock et al., 2018***). Our observations would fit with previous suggestions that a signalling gradient, such as a Notch gradient, exists across the retina and location-dependent exposure to it is important for downstream decision-making (***Murciano et al., 2002; Del Bene et al., 2008; Hiscock et al., 2018; Aggarwal et al., 2016***). Thus, our results not only have important implications for understanding the organisation of developing vertebrate tissues, but may also provide a starting point for further exploration of the connection between variability in nuclear positions and cell fate decision making in neuroepithelia.

## Methods and Materials

### Animals and Transgenic Lines

All animal work was approved by Local Ethical Review Committee of the University of Cambridge and performed in accordance with a Home Office project license PL80/2198. All Zebrafish were maintained and bred at 26.5 °C. All embryos were incubated at 28.5 °C before imaging sessions. At 10 hours post fertilization (hpf), 0.003% phenylthiourea (PTU) (sigma) was added to the medium to stop pigmentation in the eye.

### Lightsheet microscopy

Images of retinal development for the main dataset were obtained using lightsheet microscopy. Double transgenic embryos, Tg(bactin2:H2B-GFP::ptf1a:DsRed) were dechorionated at 24 hpf and screened positive for the fluorescent transgenic markers prior to the imaging experiment. The embryo selected for imaging was then embedded in 0.4% low gelling temperature agarose (Type VII, Sigma-Aldrich) prepared in the imaging buffer (0.3x Daniau’s solution with 0.2% tricaine and 0.003% PTU (***Godinho, 2011***)) within an FEP tube with 25 µm thick walls (Zeus), with an eye facing the camera and the illumination light shedding from the ventral side. The tube was held in place by a custom-designed glass capillary (3 mm outer diameter, 20 mm length; Hilgenberg GmbH). The capillary itself was mounted vertically in the imaging specimen chamber filled with the imaging buffer. To ensure normal development, a perfusion system was used to pump warm water into the specimen chamber, maintaining a constant temperature of 28.5 °C at the location of the specimen.

Time-lapse recording of retinal development was performed using a SiMView light-sheet microscope (***Tomer et al., 2012***) with one illumination and one detection arm. Lasers were focused by Nikon 10x/0.3 NA water immersion objectives. Images were acquired with Nikon 40x/0.8 NA water immersion objective and Hamamatsu Ocra Flash 4.0 sCMOS camera. GFP was excited with scanned light sheets using a 488 nm laser, and detected through a 525/50 nm band pass detection filter (Semrock). Image stacks were acquired with confocal slit detection (***Baumgart and Kubitscheck, 2012***) with exposure time of 10 ms per frame, and the sample was moved in 0.812 µm steps along the axial direction. For each time point, two 330 x 330 x 250 µm^3^ image stacks with a 40 µm horizontal offset were acquired to ensure the coverage of the entire retina. The images were acquired every 2 min from 30 hpf to 72 hpf. The position of the sample was manually adjusted during imaging to compensate for drift. The two image stacks in the same time point were fused together to keep the combined image with the best resolution. An algorithm based on phase correlation was subsequently used to estimate and correct for the sample drift over time. The processing pipeline was implemented with MATLAB (MathWorks).

### Two photon microscopy

Images for the repetition dataset and all other conditions were obtained using a TriM Scope II 2-photon microscope (LaVision BioTec). A previously established Tg(H2B-GFP) line, generated by injecting a DNA construct of H2B-GFP driven from the actin promoter (***He et al., 2012***), was used for all these experiments. Embryos were dechorionated and screened for expression of GFP at 24 hpf. An embryo was then embedded in 0.9% UltraPure low melting point agarose (Invitrogen) prepared in E3 medium containing 0.003% PTU and 0.2% tricaine. The agarose and embryo were placed laterally within a 3D printed half cylinder of transparent ABS plastic, 0.8 mm in diameter, attached to the bottom of a petri dish, such that one eye faced the detection lens of the microscope. The petri dish was then filled with an incubation solution of E3 medium, PTU, and tricaine in the same concentrations as above. For the experiment involving cell cycle arrest, hydroxyurea and aphidicolin (Abcam) were added to the incubation solution right before imaging, to a final concentration of 20 mM and 150 µM, respectively. The imaging chamber was maintained at a temperature of 25 °C, 28.5 °C, or 32 °C, as required, using a precision air heater (The Cube, Life Imaging Services).

Green fluorescence was excited using an Insight DeepSee laser (Spectra-Physics) at 927 nm. The emission of the fluorophore was detected through an Olympus 25x/1.05 NA water immersion objective, and all the signal within the visible spectrum was recorded by a sensitive GaAsP detector. Image stacks with step size of 1 µm were acquired with exposure time of 1.35 ms per line averaged over two scans. The images were recorded every 2 min for 10-15 hours starting at 26-28 hpf. The same post processing procedure for data compression and drift correction was used on these raw images as on those from lightsheet imaging.

### Obtaining experimental input values for the model

The radial coordinates *r*_*n*_ of nuclei were calculated by subtracting *l*_*n*_ from *a*, wherein *l*_*n*_ is the distance from the center of a nucleus *n* to the apical surface and *a* is the distance from the center of the lens to the apical surface. We estimated a total uncertainty of Δ*r* = ±3 µm for each single distance measurement of *r*_*n*_. This value is a result of uncertainty in detecting the center of the nucleus and in establishing the position of the apical surface.

Because each nuclear position has an error bar Δ1*r*, binning the data leads to an uncertainty in the bin count. In order to calculate this uncertainty, we considered the probability distribution of a nucleus’ position. In the simplest case, this probability is uniform within the width of the positional error bar and zero elsewhere. The probability, *p*_*n,*bin_, of finding a given nucleus *n* within a given bin, is proportional to the size of the overlap of probability distribution and bin. It follows that the expectation value for the number of nuclei within a bin is given as 𝔼 (*N*_bin_) = ∑_*n*_ *p*_*n,*bin._. Correspondingly, Var(*N*_bin_) = ∑_*n*_ *p*_n,bin_(1–*p*_*n,*bin._) is the variance of the number of nuclei within this bin. Thus, the error bar of the bin count is 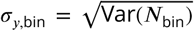. The nuclear distribution profile *N*(*r, t*) is not expected to be uniform or linear, therefore the expectation value 𝔼(*N*_bin_) does not correspond to the number of nuclei at the center of the bin. Since the position of the expectation value is unknown *a priori*, it is still plotted at the center of the bin with an error bar denoting its positional uncertainty. Here we assume this error bar to be the square-root of the bin size 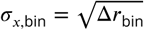.

In order to obtain the experimental nuclear concentration profile *c*(*r, t*), and its error bars, from the distribution of nuclei *N*(*r, t*), the volume of the retina also has to be taken into account, since *c* = *N*/*V*. The total retinal volume within which nuclei tracking took place was estimated directly from the microscopy images. To this end, we outlined the area of observation in each image slice using the Fiji software and multiplied this area with the distance between successive images. Given the total volume, *V*_total_, we proceeded to calculate the volume per bin, which depends on the radii at the inner and outer bin surfaces. In general, the volume of part of a sphere, e.g. a spherical sector, is given as 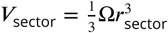, where Ω denotes the solid angle. Knowing the apical and basal tissue radii, *r* = *a* and *r* = *b*, one can thus calculate Ω as Ω = 3*V*_total_/(*a*^3^ - *b*^3^). This gives the volume of each bin as 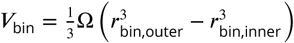, where *r*_bin,outer_ and *r*_bin,inner_ denote the outer and inner radii of a bin, respectively. Similarly, we calculated the effective surface area *S* through which the influx of nuclei occurs (see Equation 3) from the solid angle Q. This surface area is simply given as *s* = 𝔼*a*^2^.

To retrieve the average cell cycle time *T*_*P*_ for each of the data sets, we used two different approaches. In the case of the main data set, sufficient number of nuclear tracks consisting of a whole cell cycle were present. Thus we directly calculated the average cell cycle duration from these tracks. For the other datasets, we make use of the fact that the number of nuclei follows an exponential growth law depending on *T*_*P*_ (see Equation 2). Knowing the initial number of tracked nuclei *N*_0_ for each data set, we obtained *T*_*P*_ from fitting the following equation to the number of nuclei as a function of time in a log-lin plot: ln *N*(*t*) = ln *N*_0_ + *t*/*r* = ln *N*_0_ + (ln 2/*T*_*P*_)*t*. Then *T*_*P*_ was deduced from the slope of this fit.

In order to determine the maximum nuclear concentration *c*_max_ for the nonlinear model, we first randomly selected 100 nuclei from our dataset of tracked nuclei and measured the size of their longest diameter in both XY and YZ planes. From these measurements we established that the size of the principal semi-axis of each nucleus is likely to lie in the range of about 3 µm to 5 µm, where the nuclear shape is regarded to be ellipsoidal. This led to the range of possible maximum concentrations *c*_max_, although we did not measure the precise nuclear volume. The lower limit for the nuclear volume is set by the volume of a sphere of radius 3 µm, the upper limit by a sphere of radius 5 µm. Taking into account the maximum possible packing density of nuclei, which for aligned ellipsoids is the same as that of spheres (***Donev et al., 2004***), 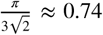, we obtained a range of 1.41 × 10^−3^ µm^−3^*c*_max_ 6.55 × 10^−3^ µm^−3^.

### Obtaining the initial condition

We determined the prefactors *h*_*i*_ from the experimental nuclear distribution at the start of the experiment, *c*_exp_(*ξ,* 0). For convenience, we chose to determine first 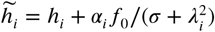 and then obtained *h*_*i*_ by subtracting 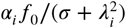 from the results. The 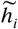 can be calculated from the data, using Equation 6 for *s* = 0, as

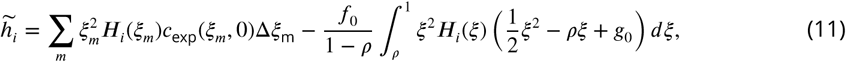

where *m* denotes the *m*-th binned data point, *ξ*_*m*_ its position and Δ*ξ*_m_ the width of bin *m*. As in Equation 6, the index *i* denotes the *i*-th eigenfunction or -mode.

### The concentration profile in the nonlinear model

The non-linear concentration profile was determined numerically from the same initial condition as used for the linear model, Equation 6, at *s* = 0 with 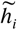 as in Equation 11. Time evolution of the initial condition, according to Equation 9, was performed using the pdepe solver in MATLAB.

### Fitting the model

The range of sizes of the nuclear principal semi-axes was used to determine the range of data to be included in our fits. Any data closer than 3 µm to 5 µm from the apical or basal tissue surfaces was not taken into account for fitting because the center of a nucleus cannot be any closer to a surface than the nuclear radius. Thus, all data collection very close to the apical or basal tissue surfaces must have been due to the above mentioned measurement uncertainties Δ*r*.

In principle, the full solution for *c*(*ξ, s*) is composed of infinitely many modes. However, in practice, we truncated this series and only included the first 8 modes in our fits. This is due to the fact that we have a finite set of data points, so adding too many modes could lead to over-fitting. Fits with a wide range of numbers of modes were found to result in the same optimal *D*-values.

For fitting, we first rescaled the data in accordance with the non-dimensionalisation of the theoretical variables *r* and *t* (see Equation 5). Thus we obtain *c*_exp_(*ξ, s*) from *c*_exp_(*r, t*). Then both models were fitted to the experimental data using a minimal-*χ*^2^ approach. The goodness of fit parameter 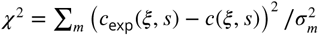 where **∑**_*m*_ denotes the summation over all bins *m*. Since binning resulted in uncertainties *σ*_*y,*bin_ and *σ*_*x,*bin_ in the *y*- and *x*-directions, both had to be taken into account when calculating *σ*_*m*_ and *χ*^2^. The combined contribution of *x*- and *y*- uncertainties is: 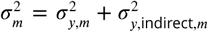 with 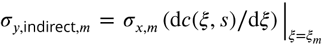 (***Bevington and Robinson, 2003***). In our fits, the value *χ*^2^ was calculated for a large range of possible diffusion constants *D*, from *D* = 0.01 µm^2^/min to *D* = 10 µm^2^/min. By finding the value of *D* for which *χ*^2^ became minimal for a given data set and time point, we established our optimal fit.

The minimal-*χ*^2^ approach furthermore enabled us to determine the optimal binning width Δ*r*_bin_ or Δ*ξ*_bin_ and width of data exclusion for the fits. In order to do so, fits of the normal data set were performed for different data binning widths and exclusion sizes of 3 µm to 5 µm. For each of these fits the *χ*^2^-value and the number of degrees of freedom *v*, i.e. the number of data points minus the number of free fit parameters (here number of data points minus 1), were registered. From *χ*^2^ and *v* we calculated the reduced *χ*^2^ value, 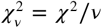 (***Bevington and Robinson, 2003***). Using *v* and 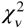, the probability *P*_*x*_ (*χ*^2^; *v*) of exceeding *χ* for a given fit can be estimated, which should be approximately 0.5 (***Bevington and Robinson, 2003***). Therefore, we found our optimal data binning width of 3 µm to 4 µm as the width that resulted in a *P*_*x*_ (*χ*^2^; *v*) as close to 0.5 as possible for all the different time points when fitting the nonlinear model. The exact choice of exclusion width was found not to influence the fitting result for the nonlinear model.

In addition to finding the optimal *D*-value for individual time points, we also modified the minimal-*χ*^2^ routine to find the value of *D* that fits a whole data set (i.e. all time points simultaneously) in the best possible way. In order to do so, we summed the *χ*^2^-values obtained for each *D* over all time points, in this way producing a ∑_*t*_ *χ*^2^(*D*)-curve. The minimum of this curve indicates *D*\* for the whole time series. Furthermore, dividing Σ_*t*_ *𝒳*^2^(*D*) by the number of time points included in the optimization yields an average *𝒳*^2^- and reduced *𝒳*^2^-value corresponding to this *D*\*. In addition, the width of this time averaged curve at 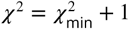 indicates the standard deviation of the optimal *D*-value, *σ*_*D*_. By approximating the minimum with a quadratic curve, we obtain an estimate for this standard deviation as 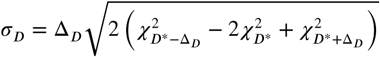 (***Bevington and Robinson, 2003***) where Δ_*D*_ is the step size between individual fitted *D*-values, here δ_*D*_ = 0.01 µm^2^/min. Lastly, based on the average reduced *𝒳*^2^-values, we also compared several *c*_max_-values for each data set to find the fit with probability *P*_*𝒳*_ (*𝒳*^2^; *v*) the closest to 0.5 in each case.

All fits were performed using custom MATLAB routines.

### *t*-tests

To compare results between data sets, the values *D*\* and corresponding *σ*_*D*_ from the overall fits were considered. It should be noted that these values were not obtained by averaging several data sets of the same experimental condition but instead each value results from one data set only. However, the sample size for each data set was set to 100 because 100 time points were taken into account for each overall optimization. These time points might not be completely uncorrelated, limiting the predictive power of the *t*-test. Two sided tests, specifically unequal variances *t*-test, also known as Welch’s *t*-test, (***Precht and Kraft, 2015***), were performed in order to determine whether samples differ significantly from each other.

## Acknowledgments

AH and REG would like to thank Oliver Y. Feng, Timothy J. Pedley, Michael E. Cates and Salvatore Torquato for helpful advice and input. This work was supported by the Cambridge Wellcome Trust PhD Programme in Developmental Biology, the Cambridge Commonwealth, European and International Trust, and Natural Sciences and Engineering Research Council of Canada (AA); Established Career Fellowship EP/M017982/1 from the Engineering and Physical Sciences Research Council (REG); and Wellcome Trust Investigator Award (SIA 100329/Z/12/Z) (WAH).

## Appendix

### Full solution of the linear diffusion equation

After rescaling space and time as in Equation 5 and introducing *ρ* = *a*/*b* < 1, Equation 1 and the boundary conditions 3 and 4 read

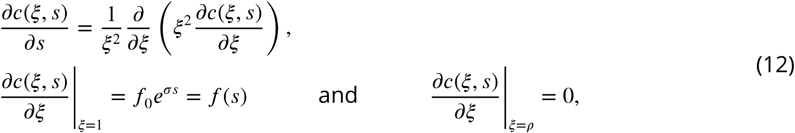

where we have defined *f*_0_ = *aN*_0_/*DSτ* and *σ* = *a*^2^/*Dτ*. We transform this homogeneous differential equation with inhomogeneous boundary conditions into the problem of solving an inhomogeneous differential equation with homogeneous boundary conditions by writing *c*(*ξ, s*) as a sum of two contributions,

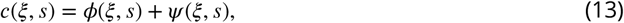

where we require *ϕ*(*ξ, s*) to satisfy the inhomogeneous boundary conditions

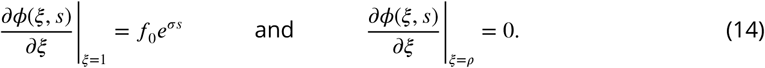

These conditions are satisfied if *ϕ*(*ξ, s*) has the form

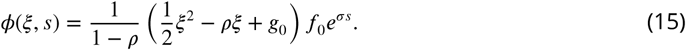

where *g*_0_ is a constant of integration to be determined later. The remaining problem to solve for *ψ*(*ξ, s*) is

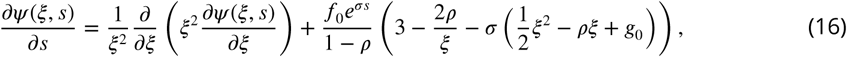

with homogeneous boundary conditions

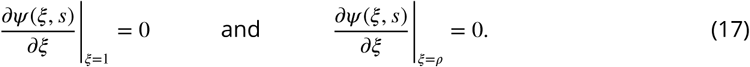

We can further write *ψ*(*ξ, s*) as the sum of two contributions,

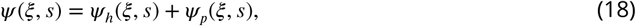

where *ψ* _*h*_ is the general solution of the homogeneous problem

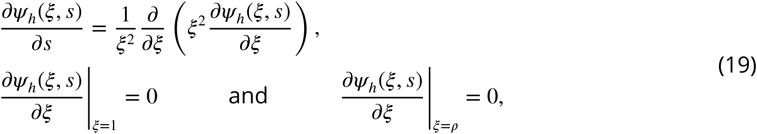

and *ψ* _*p*_ is a particular solution of the full inhomogeneous problem 17. The full solution of the homogeneous problem is given as a series of linearly independent eigenfunctions, each of the form

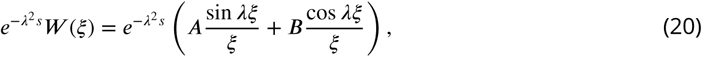

where the eigenvalues *λ* can be found from simultaneous solution of the boundary conditions,

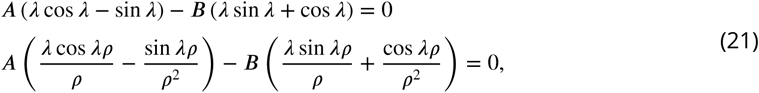

which yields the transcendental relation

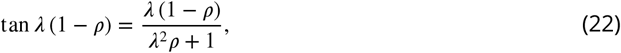

for which each eigenvalue *λ* _*i*_ is a solution corresponding to one of the linearly independent eigen-functions (only *λ*_*i*_ > 0 need to be taken into account). We can further deduce from the Equation 21 that *B*_*i*_ = *β*_*i*_*A*_*i*_, where

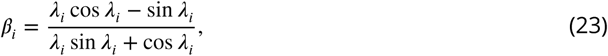

and we normalize the obtained expression for *W*_*i*_(*ξ*) from Equation 20

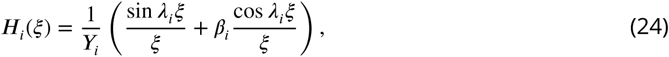

with

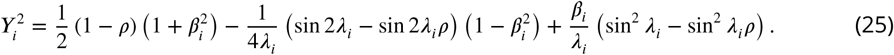

Thus, the homogeneous solution *ψ*_*h*_ is

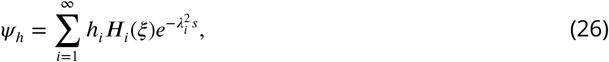

with prefactors *h*_*i*_ to be determined from the initial condition.

In order to find a particular solution of the inhomogeneous problem, we first rewrite 17 as

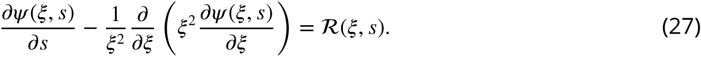

Now, we express ℛ(*ξ, s*), as well as the unknown inhomogeneous solution *ψ*_*i*_ (*ξ, s*) in terms of the normalized eigenfunctions *H*(*ξ, s*) of the homogeneous problem,

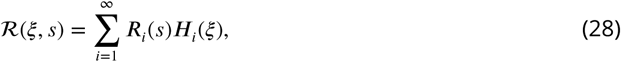

and

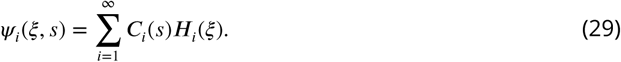

Substituting these forms into 27, and noting that each term in the series must vanish separately we obtain

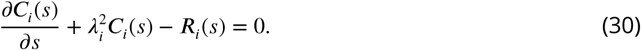

From the form of ℛ(*ξ, s*) it follows that *R*_*i*_(*s*) = *α*_*i*_*f*_0_ *e*^*σs*^ with some purely numerical prefactors *α*_*i*_, so we expect *C*_*i*_(*s*) ∝ *p*_*i*_*e*^*σs*^ and find

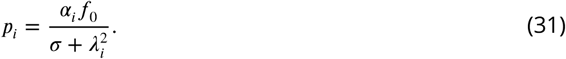

Finally, we determine the *α*_*i*_ by reconsidering Equation 28. We multiply both sides by *ξ*^2^*H*_*j*_(*ξ*), where *H*_*j*_(*ξ*) is one specific but arbitrary eigenfunction of the homogeneous problem, and then integrate over the whole volume *V*. By the orthogonormality of these eigenfunctions we obtain

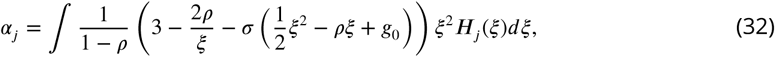

and all the *α* _*i*_ can be calculated explicitely. Thus, the full solution of the linear problem is

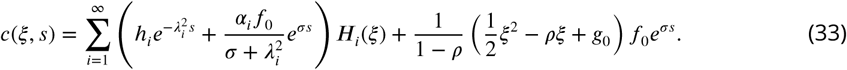

The constant *g*_0_ can now be calculated from the requirement that *∫ c*(*ξ, s* = 0)*dV* = *N*_0_. Here we make use of the fact that *∫ H*_*i*_(*ξ*)*ξ* ^2^*d ξ* = 0 if *λ*_*i*_ satisfies Equation 22, thus

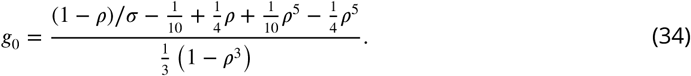

